# Proteonano™: a novel deep proteomics platform with 1000-plex profiling capacity and picogram sensitivity and its application in diabetic kidney disease

**DOI:** 10.1101/2023.09.12.556305

**Authors:** Ban Zhao, Xuechun Gao, Xiehua Ouyang, Jiakai Fang, Zihao Deng, Hao Wu, Yonghui Mao

## Abstract

The development of blood-based multi-biomarker panels for screening diabetic patients, and as an easy-to-access tool for identifying individuals at greatest risk of developing diabetic kidney disease (DKD) and its progression, is essential. However, conventional blood biomarker-based methodologies (e.g. clinical tests and ELISA) are unable to predict DKD progression with high sensitivity and specificity. To overcome these challenges, we developed a deep, untargeted plasma proteome profiling technology (Proteonano™ platform) to identify potential multiple protein biomarkers involved in DKD progression. The Proteonano™ technology is an affinity selective mass spectrometric platform that comprises nanoparticle-based affinity binders (nanobinders) for low abundant protein enrichment, automated workflow for parallel sample preparation, and machine learning empowered bioinformatic software for data analysis.

Using the Proteonano™ platform, we performed untargeted proteomics on 75 subjects (DKD progressors, n = 30; DKD non-progressors, n = 45) and identified an average of 953 ± 80 (AVG ± SD) protein groups, with a wide dynamic range of 8 orders of magnitude (with the lowest concentration down to 3.00 pg/mL). Among these, 38 proteins were differentially expressed between DKD progressors relative to non-progressors, and the predictive power for these proteins were assessed. Further, we performed random forest and LASSO analyses for additional variable selection. Variables selected by these approaches were assessed by Akaike information criterion method followed by ROC analysis, which identified a combination of multiple proteins (including VWF, PTGDS, B2M, BT3A2, and LCAT) that showed excellent predictive power over current methods, with an area under the curve value up to 0.97. Some of these plasma proteins are not previously recognized in the context of DKD progression, suggesting they are novel biomarkers. Our studies pave the way to develop multi-biomarker panels for DKD progression management. This study suggests that the Proteonano™ technology platform reported here can be employed as an established workflow enabling untargeted deep proteomic analysis to identify highly discriminative biomarkers for precise medicine.

## INTRODUCTION

Diabetes is a global health challenge that affects about 500 million people around the world [1]. One of the major long-term complications for diabetes is diabetic kidney disease (DKD) [2, 3]. About 30 to 40 % of type I and type II diabetes patients develop DKD overtime [2, 3]. DKD is believed as a leading cause of end-stage kidney disease (ESRD), and about 40 % ESRD patients that requires kidney transplant suffer from mild to severe DKD [2, 3]. So far, DKD is considered as a microvascular disorder, and the entry of excess glucose to endothelial and other cells are viewed as a major driving force for disease development [2, 3]. However, only a fraction of diabetes patients progress into DKD, and within the DKD population, the pathology of disease progression is also heterogenous [2, 3]. Thus, it is imperative to identify those patients who have higher likelihood to developing DKD and higher risk for DKD progression at early stages. Easy-to access diagnostic panels that have highly sensitive and specific prognostic biomarkers for DKD progression will be beneficial to improve life of millions of patients [4–7].

Urine albumin and estimated glomerular filtration rate (eGFR) are two common biomarkers used to diagnose and predict DKD progression [4–7]. However, the specificity and sensitivity of these two biomarkers are relatively low. For example, a fraction of DKD with type II diabetes do not develop albuminuria, and albuminuria and loss of eGFR are decreased in other form of kidney diseases [4–7]. In the case of eGFR, it does not always reflect the severity of kidney damage, as increase in single nephron glomerular filtration rate may compensate for the loss of functional nephron [4–7]. Thus, patients with the same eGFR may have drastically different prognosis [4–7].

Thus, it is critical to develop novel biomarkers for DKD diagnosis and prognosis. Based on current knowledge of DKD pathology, a large number of substances that are associated with inflammation, endothelial dysfunction, fibrosis, and renal tubule damage have been screened for their potential serving as biomarkers. Despite of the plethora of candidate biomarkers, few can improve the predictive power of existing models that incorporate clinical variables, urine albumin, and eGFR [4–8].

Recently, high resolution mass spectrometric proteomics approaches were employed to discover novel biomarkers for DKD management. One of the urine proteomics studies using chronic kidney disease (CKD) identified 273 urinary peptides that can be combined into a single classifier to predict eGFR loss [9–11]. This assay has been validated in larger cohorts and is commercially available [8]. Moreover, with the production of multi-protein panels, a sizable number of studies using protein binding aptamers and proximity extension assays have unveiled novel candidate biomarkers for kidney diseases [12–15].

Despite these progresses, these assays have their own limitations. For example, targeted panels are limited by the need to establishing individual antibodies to detect each protein, and urine proteomics is limited by proteins and peptides that present in urine samples. Developing blood based untargeted liquid chromatography-tandem mass spectrometry (LC-MS/MS) methods is more powerful approach to discover novel plasma biomarkers; however, current methodologies based on LC-MS/MS are largely restricted by complicated sample preparation, low throughput, and relative high detecting threshold.

To circumvent these limitations, we developed a deep, untargeted plasma proteome profiling technology (Proteonano™ platform) to identify multiple protein biomarkers involved in the pathology of DKD progression. Our proof-of-concept prospective study using 75 DKD patient serum samples was able to identify 38 significantly altered proteins that are associated with DKD progression. Using multivariate analysis and machine learning approaches, we identified multivariable combinations with significant sensitivity and specificity (AUC up to 0.97). These results provide previously underappreciated potential prognostic biomarkers for DKD, which may also provide novel mechanistic insights for DKD development.

## RESULTS

### Patient characteristics

75 patients with primary clinical diagnosis of diabetic kidney disease, but not other kidney disease and hypertension were serially enrolled from Beijing Hospital Nephrology Clinic between 2012 and 2019 with ethical committee approval. At the time of enrollment, serum creatinine was determined and serum samples were banked for further use. Patients were followed up for up to five years, and were stratified by serum creatinine concentration change. Patients with urine creatinine increase more than 50 mmol/L (progressor group, P) and patients with urine creatinine increase less than 50 mmol/L (non-progressor group, NP) were compared. Except serum creatinine, demographic parameters were similar for both groups at the time of enrollment (**TABLE 1**).

**TABLE 1.** Clinical characteristics of enrolled patients.

|  | Age (y) | Gender<br>(female) n (%) | Observation<br>Period (y) | Serum<br>Creatinine at<br>Enrollment | Serum<br>Creatinine at<br>End of<br>Observation<br>Period | Serum<br>Creatinine<br>Concentration<br>Change |
| --- | --- | --- | --- | --- | --- | --- |
| Non-progressors<br>(n=45) | 62.1 ± 1.3 | 13 (28.9%) | 3.9 ± 0.2 | 87.3 ± 4.3 | 98.3 ± 5.0 | 11.0 ± 3.1 |
| Progressors<br>(n=30) | 58.3 ± 1.9 | 7 (23.3%) | 3.6 ± 0.2 | 147.9 ± 19.9 | 415.7 ± 63.4 | 283.7 ± 47.3 |

### Untargeted deep proteomics identified proteins associated with DKD progression

To identify individual biomarkers, or combination of biomarkers that can distinguish DKD patients that progressed within up to 5 years and patients that did not progress, we took a non-targeted deep proteomics approach [16, 17].

All samples were subjected to same preprocessing by first incubating serum samples with nanobinders, followed by LC-MS/MS data acquisition, peptide mapping, and protein inference [18–20]. A total of 1,393 proteins were identified in all samples, and 953 ± 80 (AVG ± SD) protein groups were identified in each serum sample (**FIGURE 2, A**), with 469 protein groups identified in all samples. Out of all the protein groups identified, 1,185 could be matched to the human plasma proteome project (HPPP) database (**FIGURE 2, B**) [21, 22], which contains 4,608 proteins. Out of these protein groups, 1,178 of them have concentration information available. Identified protein with highest concentration in the plasma is albumin (ALBU, P02768), at reported concentration of 860,000 ng/mL, and identified protein with lowest reported plasma concentration is inositol 1,4,5-triphosphate receptor associated 2 (IRAG2, Q12912), at 3.00 pg/mL (**FIGURE 2, C**). Thus, the abundance of these proteins spans across more than eight orders of magnitude.

**FIGURE 1.**
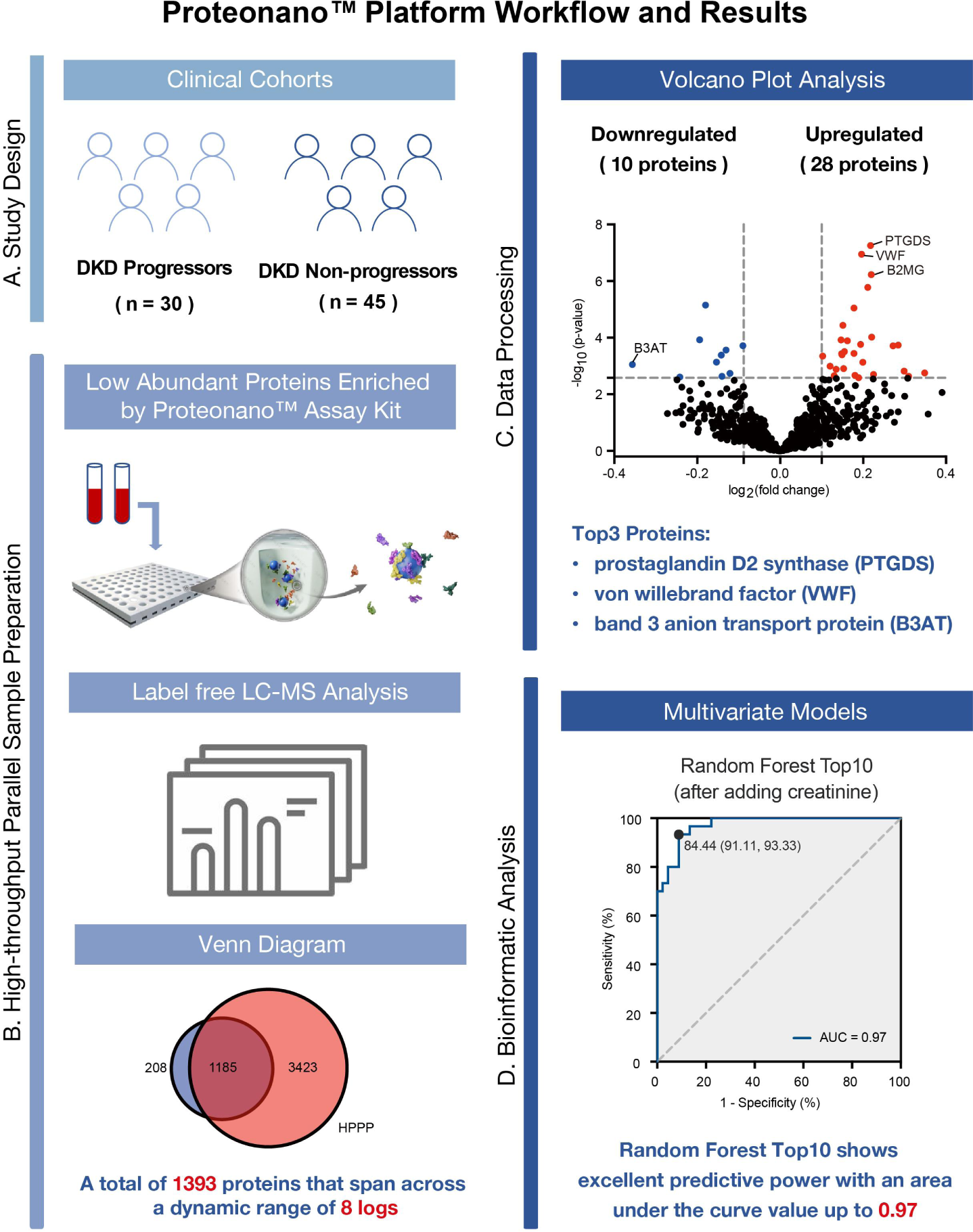
(A) Study design. (B) Proteonano ™ platform workflow for high-throughput proteomic study. (C) Volcano plot shows the top 3 differentially expressed proteins between the two cohorts. (D) Random Forest Top10 shows excellent predictive power with an AUC up to 0.97.

**FIGURE 2.**
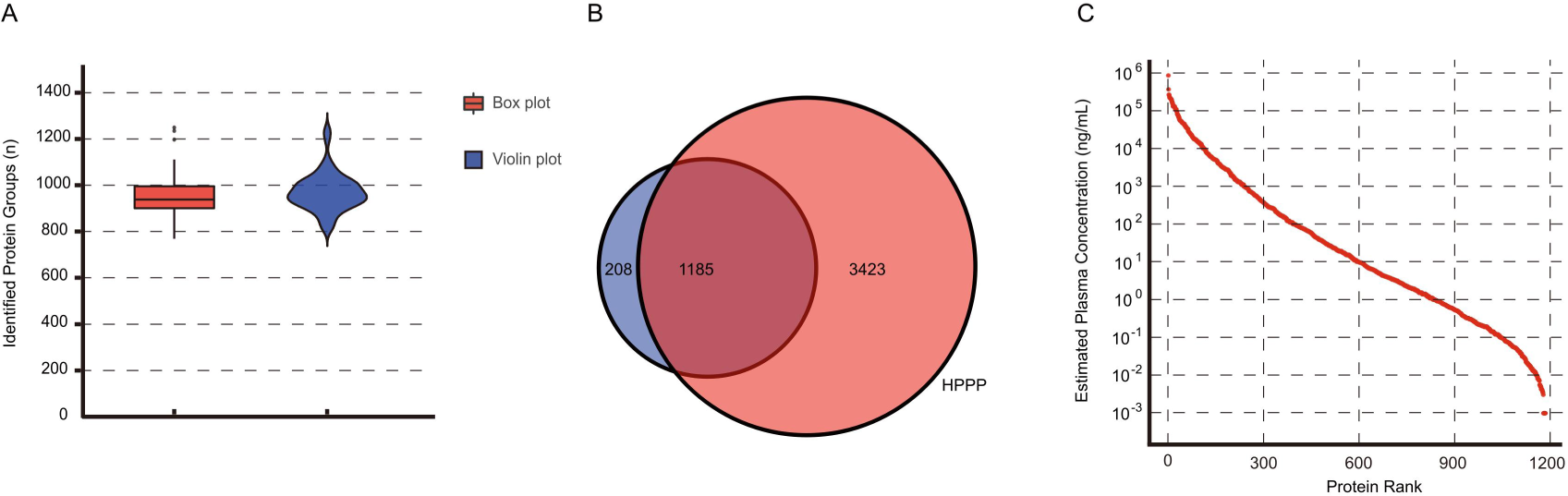
The results of Proteonano^TM^ based proteomic analysis. (A) Boxplot (red) and violin plot (blue) show the distribution identified protein groups for each sample. (B) Venn diagram demonstrates overlapped proteins between the identified ones in this study (blue) and those listed in the Human Plasma Proteome Project (HPPP, red). (C) A total of 1393 proteins that span across a dynamic range of 8 logs. Protein concentration of identified protein groups is based on plasma protein concentration reported by human plasma proteome project.

After data normalization, differential expression analysis was performed on all of the 1,393 proteins by using the Perseus software. This identified 38 proteins exhibiting significant differences between DKD patient that progressed during observation period, and patient that did not progress. This includes 10 downregulated proteins and 28 upregulated proteins in DKD progressors (**FIGURE 3, A**). Despite differential expression, odds ratio calculations suggested that most of the differentially expressed proteins we identified cannot sufficiently distinguish DKD progressors and non-progressors (**FIGURE 3, B**).

**FIGURE 3.**
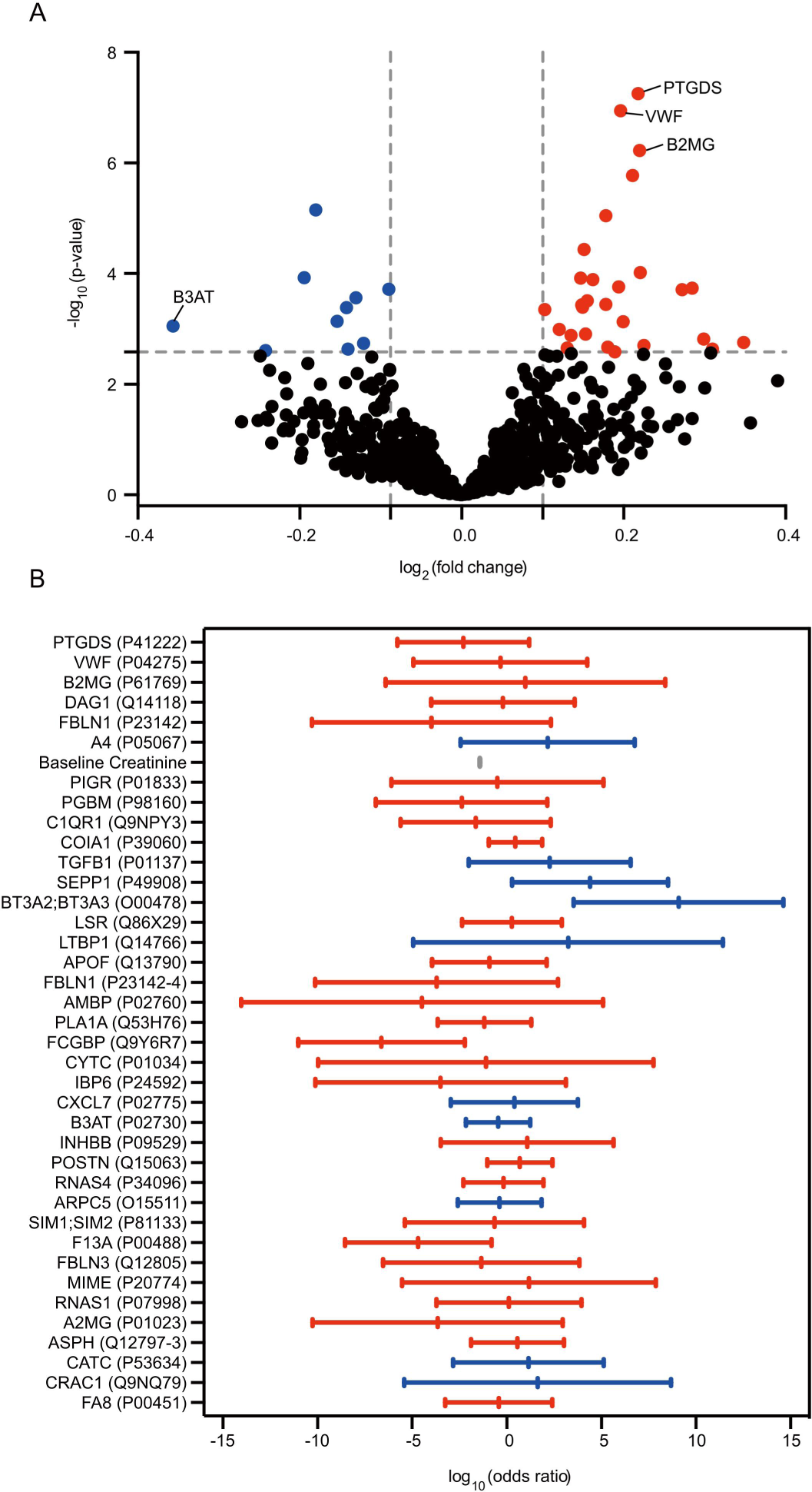
Differentially expressed proteins in DKD patients. (A) Volcano plot shows differentially expressed proteins in DKD progressors and non-progressors. X-axis shows the log_2_(fold change) of protein abundance between the two groups. Y-axis shows the -log_10_(p-value). Red dots represent 28 upregulated proteins in DKD progressors, and blue dots represent 10 downregulated proteins in DKD progressors (FDR corrected p<0.05). (B) Odds Ratio (OR) analysis of 38 candidate proteins. Log_10_(odds ratios) and 5-95 % confidence intervals are shown for each candidate protein, along with values for baseline serum creatinine. Red bars represent 28 upregulated proteins in DKD progressors, and blue bars represent 10 downregulated proteins in DKD progressors.

### Candidate biomarker proteins participate a number of cellular functions

We next probed which pathways these the identified 38 candidate protein biomarkers may participate and the relationship between these proteins. We first performed gene ontology (GO) biological process and Kyoto encyclopedia of genes and genomes (KEGG) pathway enrichment analyses (**FIGURE 4**) [23, 24]. GO analysis showed these proteins participate in 1) coagulation and hemostasis, 2) organization, assembly, regulation of the extracellular matrix, and responses to cholesterol and steroid compounds. For biological processes and pathways, these proteins involve in tissue development, maintenance, and responses to injury or external stimuli [25, 26]. For molecular functions, they are related to extracellular matrix structural components, peptidase, endopeptidase, and nucleic acid endonuclease enzyme activities, and binding functions of immunoglobulins, fibronectins, growth factors, and proteases. These proteins may function in multiple cellular components, including platelet alpha granules and their lumen, Golgi apparatus, basement membrane, and low-density lipoprotein particles. Out of these, platelet alpha granules and platelet alpha granule lumen are components in platelets responsible for coagulation and fibrinolysis, which can affect thrombus formation and bleeding-related diseases [27, 28]. Proteins in the Golgi apparatus lumen can influence protein regulation and metabolic pathways, while abnormalities in lipid metabolism may lead to cardiovascular diseases and liver dysfunction [29].

**FIGURE 4.**
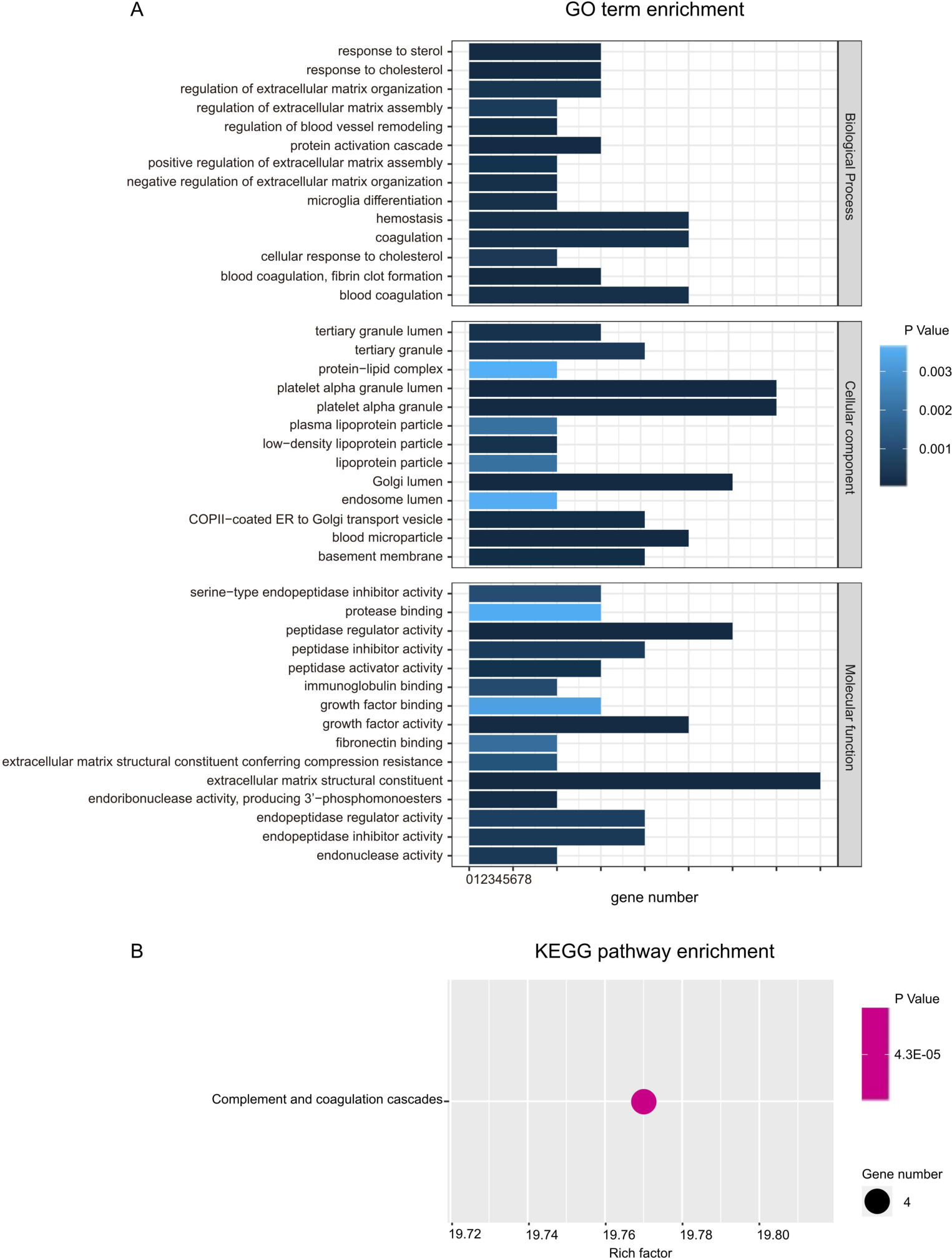
Candidate protein GO term enrichment analysis and KEGG pathway analysis. (A) GO enrichment analysis. Horizontal axis represents the number of genes involved. The depth of color corresponds significance of pathway enrichment. (B) KEGG pathway enrichment analysis. Horizontal axis represents KEGG enrichment factor; a higher enrichment factor value indicates that the abundance of differentially expressed genes in this functional category is larger compared to the overall background gene set. The size of the bubble represents number of genes in the pathway, while the color indicates the p-value.

KEGG signaling pathway analysis identified the candidate protein biomarkers may participate in complement and coagulation cascades (**FIGURE 4, B**). Four candidate marker genes were found to participate in this pathway, and this pathway participates in processes regulating cardiovascular functions [30].

STRING analysis was performed to predict functional associations among candidate proteins (**FIGURE 5**) [31]. A total of 75 functional associations were identified among candidate proteins. Within this network, B2M, PTGDS, and VWF are significant nodes, having multiple functional associations with other proteins. B2M and PTGDS are co-expressed, and their co-occurrence is well recognized [32, 33]. The relatively short link between them indicates that their predicted association is reliable. Noticeably, both B2M and PTGDS have no functional association with VWF, suggesting they may belong to two separate regulatory pathways.

**FIGURE 5.**
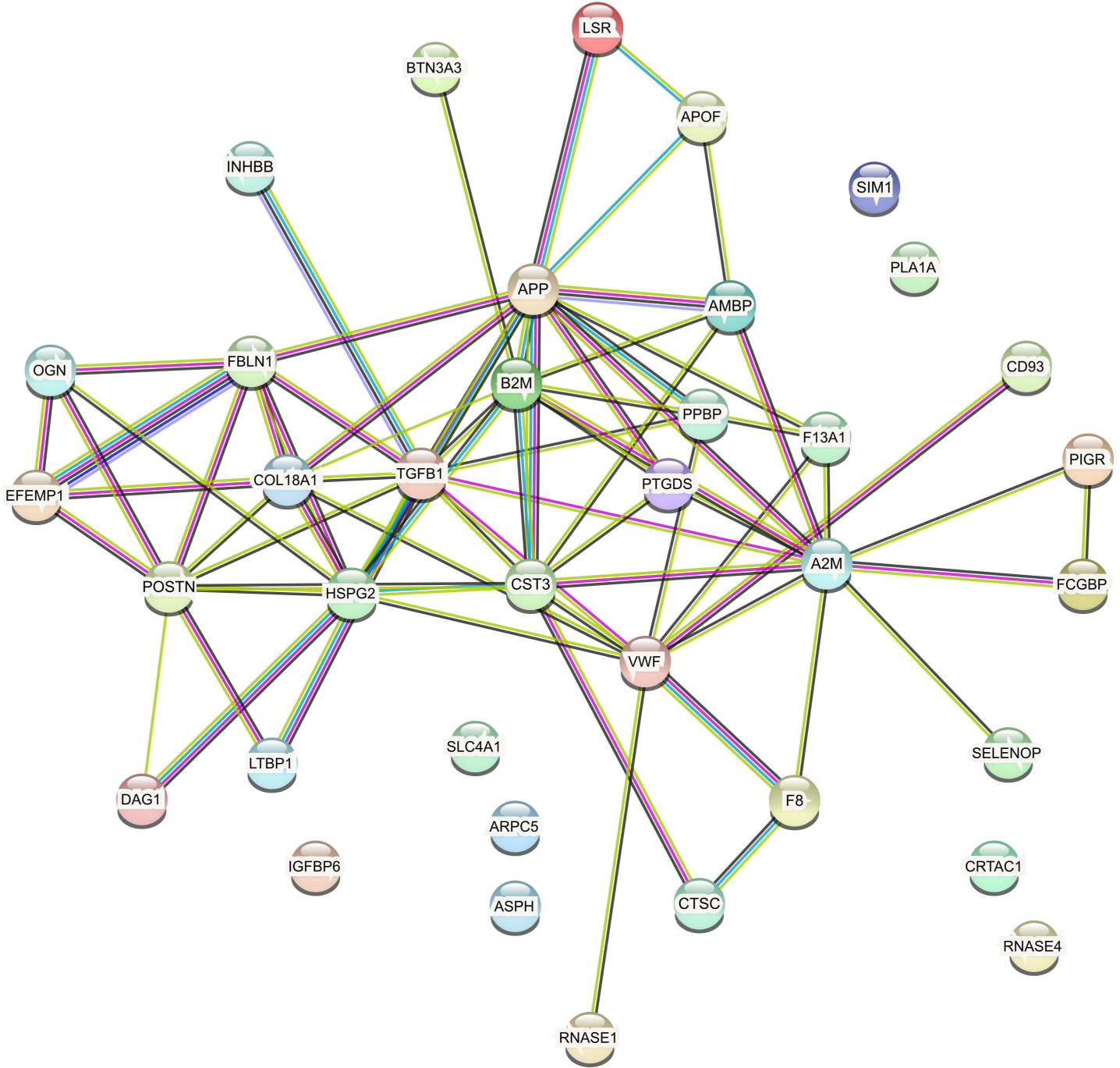
Candidate protein interaction network STRING analysis. Different line colors between proteins represent the types of associations. Blue lines correspond to the protein-protein interactions obtained from high-throughput experiments. Green lines represent interactions obtained from homology-based predictions. Red lines signify interactions derived from small-scale experiments. Purple lines indicate interactions obtained from text mining techniques. Yellow lines represent interactions predicted by gene neighborhood methods. Black lines represent interactions predicted via co-expression analysis based on similar expression patterns of the two genes across multiple conditions, suggesting that they may interact with each other. Light blue lines represent interactions predicted through fusion events. The length of lines represent the confidence of the interaction associations; shorter distances indicate more reliable predictions.

### Candidate proteins as individual predictors for DKD progression

Subsequent to identification of individual candidate proteins, we further determined their capability to differentiate between DKD progressors and non-progressors. Expression profiles for these proteins across all samples were visualized by box plots, and predictive power for each candidate protein was assessed by receiver operating characteristic (ROC) curve analysis and area under the curve (AUC) value determination (**FIGURE 6**) [34]. Notably, four proteins, namely PTGDS (P41222, AUC=0.84), VWF (P04275, AUC=0.84), B2MG (P61769, AUC=0.83), and DAG1 (Q14118, AUC=0.80) demonstrated the most robust discriminatory power, with elevated expression in DKD progressors (**FIGURE 6, C**). Pearson correlation analysis was also performed to explore the relationship among these proteins, the results were shown in heatmaps (**FIGURE 7**). Fold change analysis unveiled 11 proteins exhibiting significant differential expression above 2-fold (p < 0.05).

**FIGURE 6.**
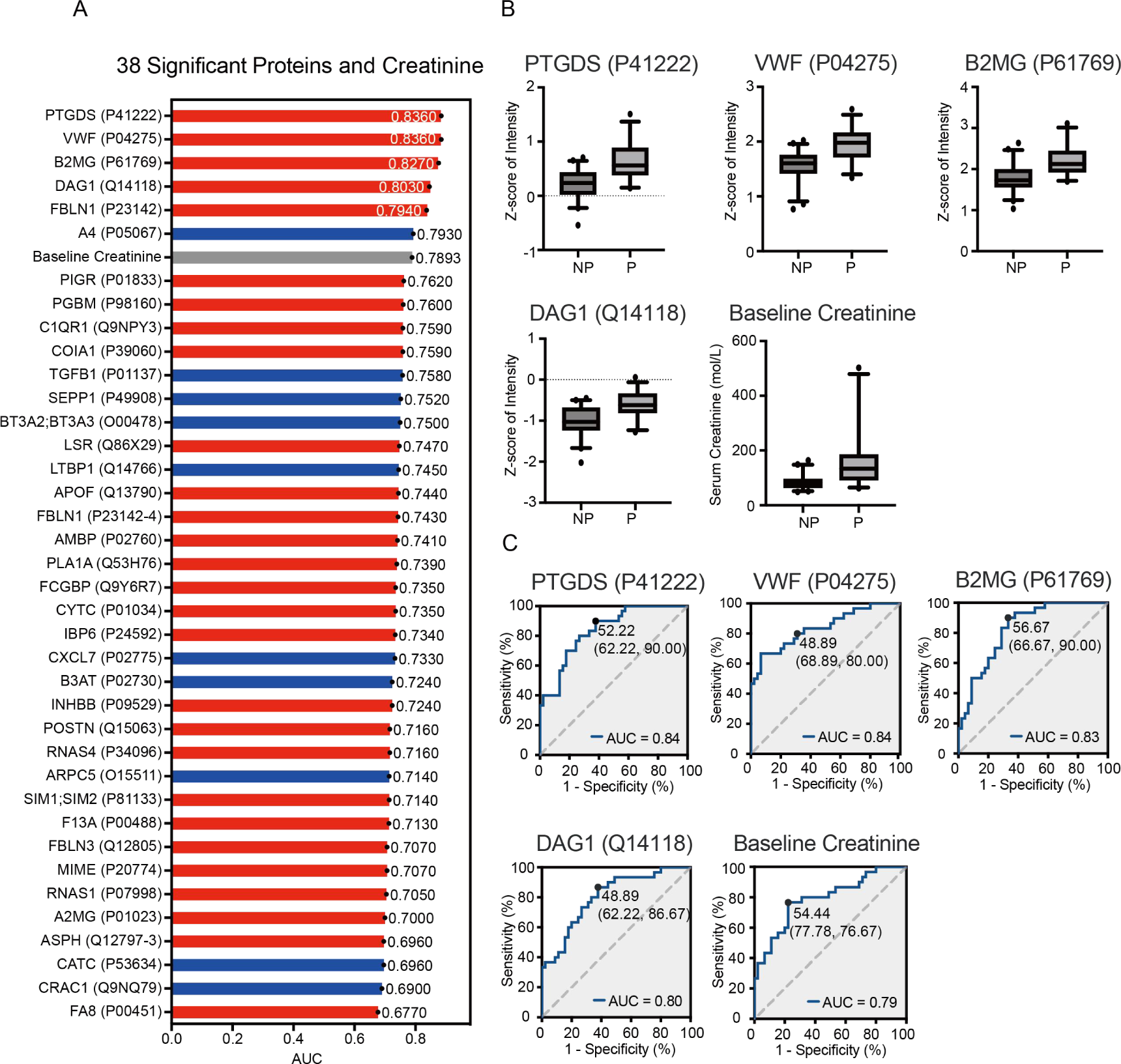
Predictive power of individual candidate proteins. (A) AUC values of ROC curves for baseline serum creatinine and 38 candidate proteins. Red bars indicate proteins upregulated in DKD progressors, blue bars indicate proteins downregulated in DKD progressors. AUC values for baseline serum creatinine are shown in the gray bars. (B) Box plots of the top 4 proteins with highest AUC values differentiating progressors and non-progressors, including PTGDS (P41222), VWF (P04275), B2MG (P61769) and DAG1 (Q14118). Expression levels are displayed as log2 transformed normalized intensities. (C) ROC curves of top 4 proteins with highest AUC values. AUC values are shown on the bottom of each panel. Youden index (specificity, sensitivity) values are also shown in each panel.

**FIGURE 7.**
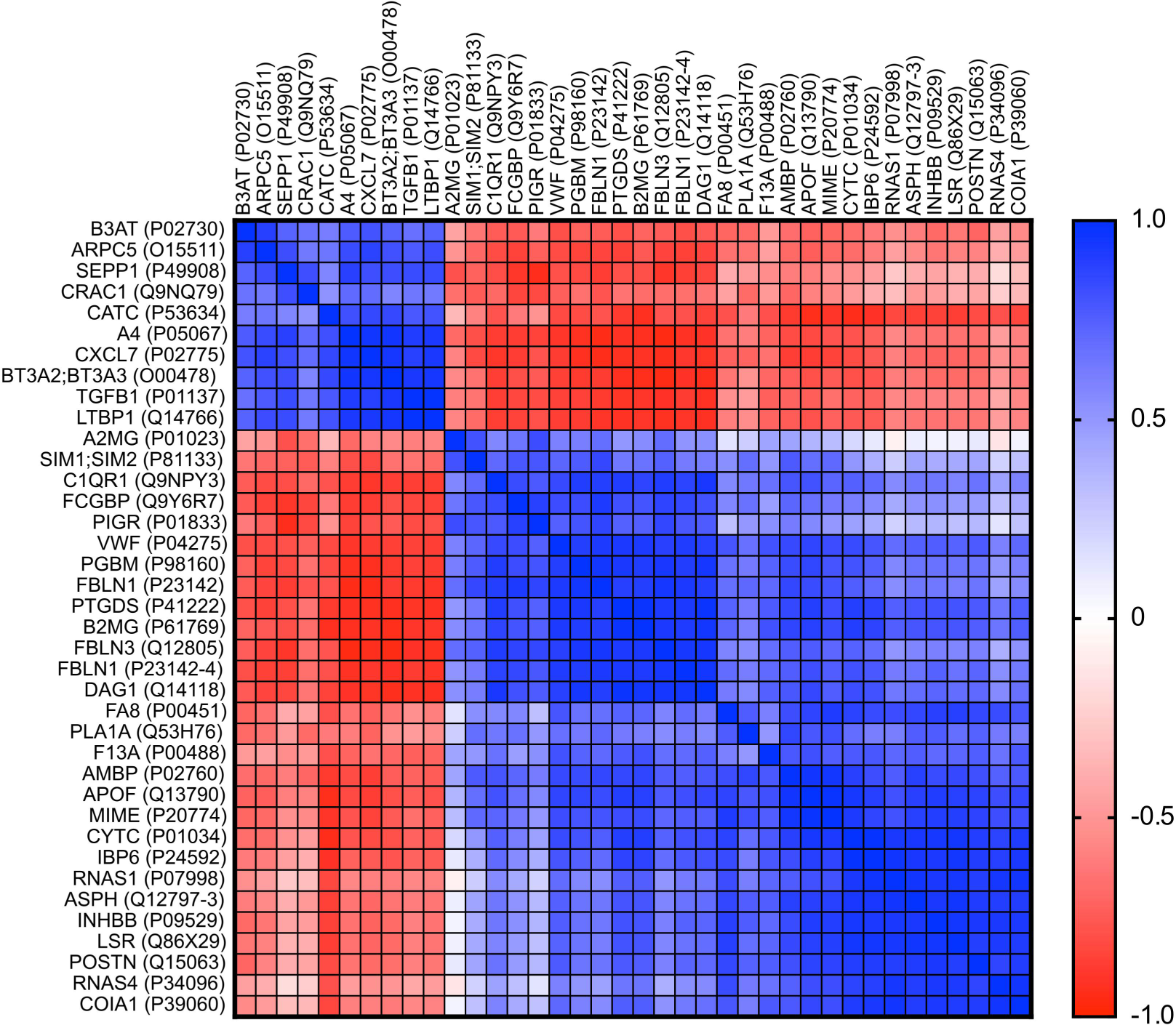
Pearson correlation analysis of the candidate proteins. Red squares indicate positive correlation while blue squares indicate negative correlation. The color intensity is proportional to correlation coefficients.

### Use of multiple candidate proteins as combined predictors for DKD progression

Although univariate ROC analysis demonstrated promising predictive power for some of the proteins identified, the specificity and sensitivity of these models remained insufficient. Thus, we investigated whether more sophisticated models using combined protein candidates would lead to an improved predictive power. To identify potential serum protein biomarker combinations, random forest and least absolute shrinkage and selection operator (LASSO) models were created using all the proteins identified in LC-MS/MS experiments [35, 36]. Top 10 proteins were identified in both analyses (**FIGURE 8, A-B**).

**FIGURE 8.**
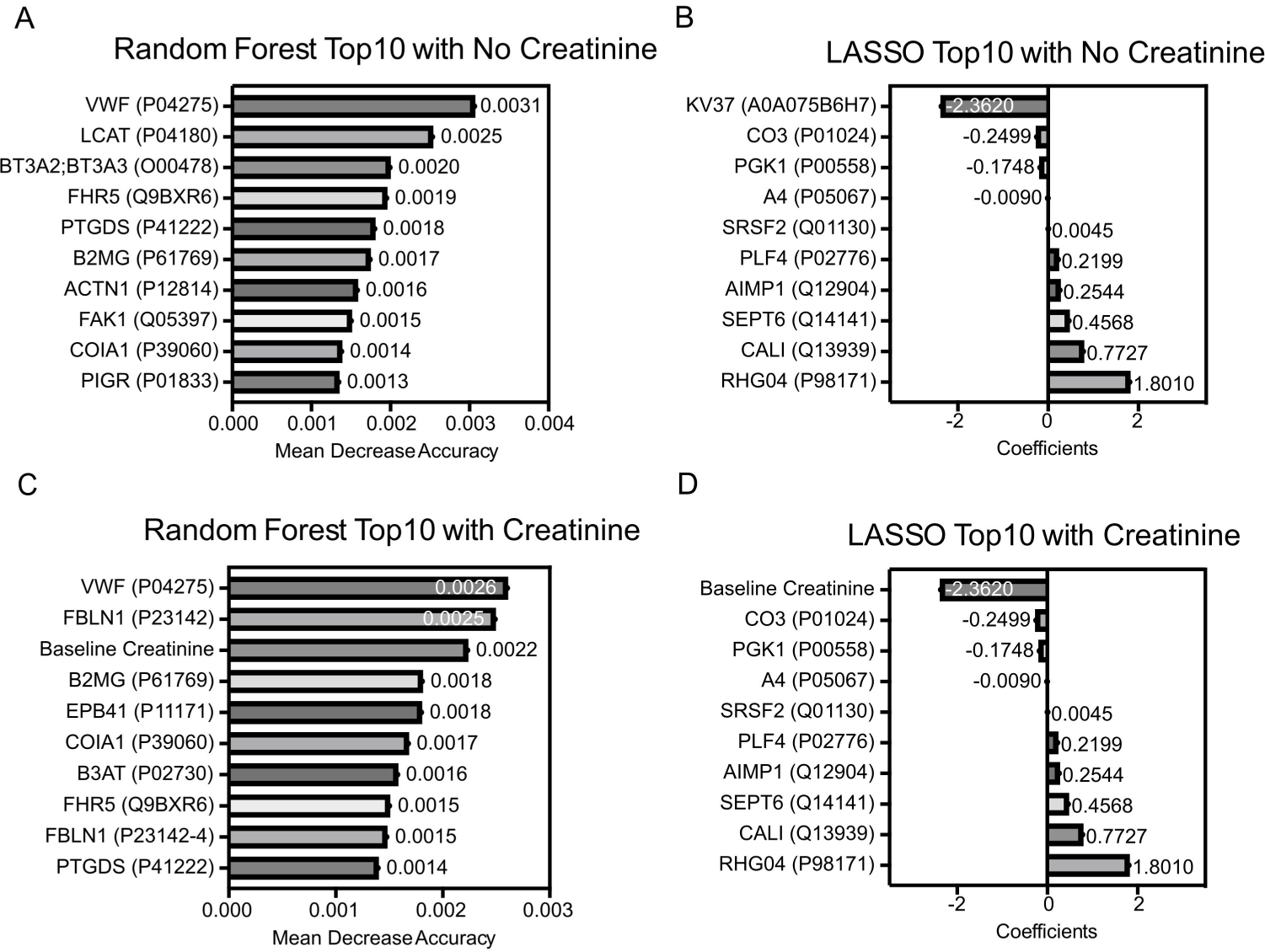
Results of the random forest and LASSO analyses. Feature selection was performed to identify key predictive proteins using random forest (A, C) and LASSO (B, D) models. (A, B) Top 10 proteins identified from random forest and LASSO analyses without addition of baseline serum creatinine as a variate. (C, D) Top 10 proteins identified from random forest and LASSO analyses with addition of baseline serum creatinine as a variate.

Integrating results from univariate (fold change and ROC), random forest, and LASSO analyses, a total of 12 multivariate models were evaluated (**TABLE 2**). Akaike information criterion (AIC) was utilized to identify the optimal model (model 1, Random Forest Top10) [37], with an AIC value of 53.46 and the highest AUC value of 0.97. The resulting 7 models with AUC values exceeding 0.90 were presented in **FIGURE 9, A**. Notably, PTGDS (P41222) had the highest AUC value in univariate analysis. It also was one of the top 5 proteins selected from random forest and LASSO analyses, suggesting it could be directly associated with DKD.

**TABLE 2.** Multivariate model selection and performance assessment. Features were selected based on random forest, LASSO, AUC, and fold change analyses. Model performance was compared based on AUC, AIC and intercept values.

| Group | Combination Name | AUC | AIC | Intercept |
| --- | --- | --- | --- | --- |
| Model 1 | Random Forest Top10 | 0.97 | 53.05 | -8.33 |
| Model 2 | LASSO Top10 | 0.85 | 90.87 | -14.14 |
| Model 3 | Random Forest Top3 | 0.93 | 58.68 | -0.42 |
| Model 4 | LASSO Top3 | 0.68 | 101.96 | 9.37 |
| Model 5 | Random Forest Top3 + LASSO Top3 | 0.95 | 57.21 | 17.56 |
| Model 6 | Random Forest Top2 + LASSO Top2 | 0.89 | 68.14 | 0.04 |
| Model 7 | Random Forest Top1 + LASSO Top1 | 0.86 | 71.80 | -0.85 |
| Model 8 | Random Forest Top3 + LASSO Top1 | 0.95 | 54.57 | 14.81 |
| Model 9 | AUC Top4 | 0.93 | 62.30 | -9.96 |
| Model 10 | Fold Change Top3 | 0.82 | 82.65 | -1.31 |
| Model 11 | AUC Top4 + Fold Change Top3 | 0.95 | 57.41 | -14.32 |
| Model 12 | AUC Top1 + Fold Change Top1 | 0.93 | 58.25 | -13.39 |

**FIGURE 9.**
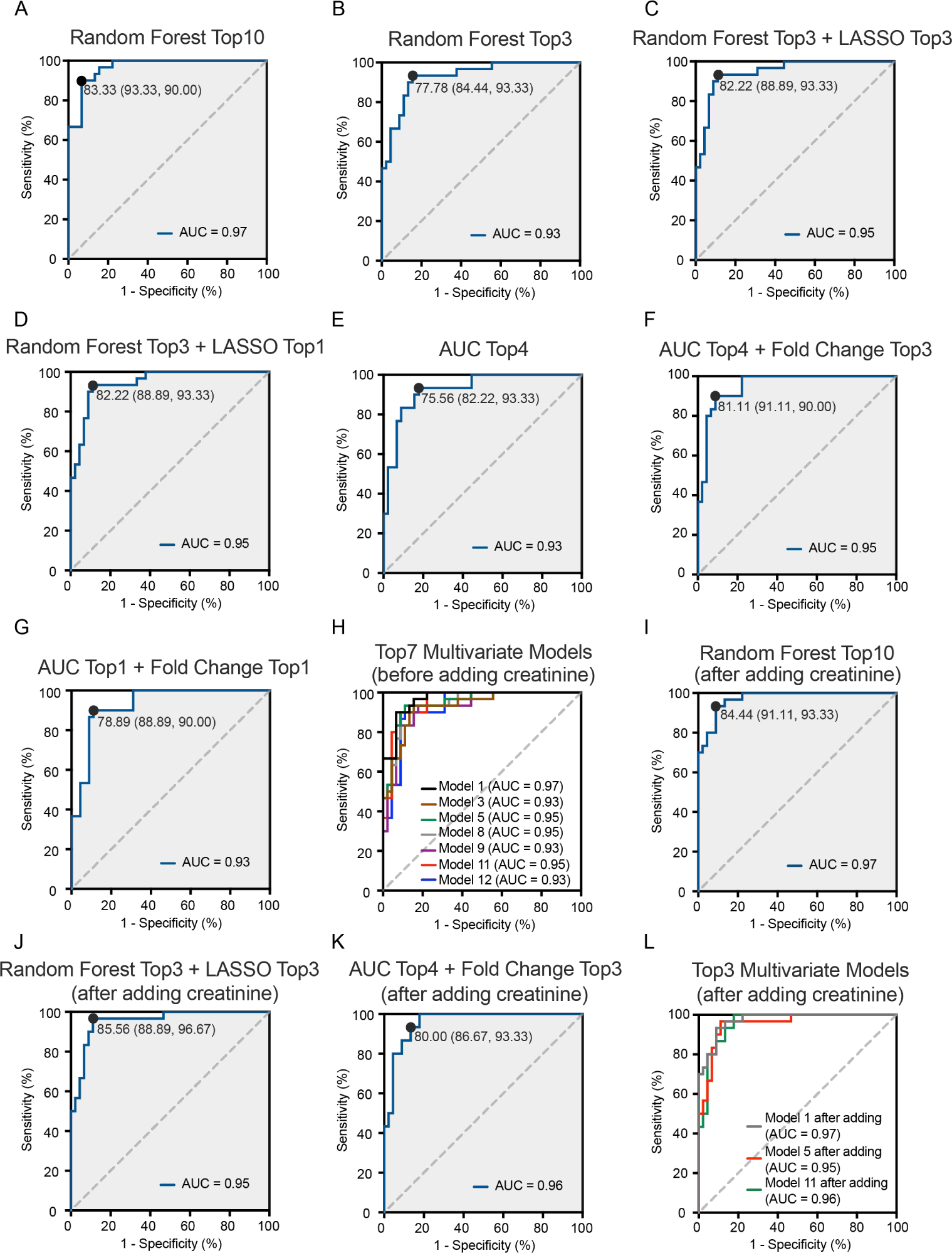
Assessment of top multivariate models. (A-G) ROC curves of top multivariate models. AUC value, Youden index (sensitivity, specificity) are indicated in each panel. (H) ROC curves of top 7 multivariate models. (I-K) RUC curves of top models with addition of baseline serum creatinine as a variate. AUC value, Youden index (sensitivity, specificity) are indicated in each panel. (L) ROC curves of top 3 multivariate models after addition of baseline serum creatinine as a variable.

Considering the potential effect of baseline serum creatinine on model selection, we included it as a variable in additional random forest and LASSO analyses (**FIGURE 8, C-D**) [38]. However, the inclusion did not lead to an improved model (data not shown). When baseline serum creatinine was added to the top 4 models with AUC values exceeding 0.95 (as shown in **FIGURE 9, A-H**), this did not improve calculated AIC and AUC numbers (**TABLE 3** and **FIGURE 9, I-L**).

**TABLE 3.** Assessment of multivariate models after addition of serum creatinine as a variate. Performance of top 3 multivariate models with and without adding baseline serum creatinine as a variate was compared. Model performance was compared based on AUC, AIC and intercept values in models with and without adding baseline serum creatinine as a variate.

| Group | Combination Name | AUC | AIC | Intercept |
| --- | --- | --- | --- | --- |
| Model 1 | Random Forest Top10 | 0.97 | 53.05 | -8.33 |
| Model 1 + Creatinine |  | 0.97 | 54.13 | -8.28 |
| Model 5 | Random Forest Top3 + LASSO Top3 | 0.95 | 57.21 | 17.56 |
| Model 5 + Creatinine |  | 0.95 | 55.78 | 16.86 |
| Model 11 | AUC Top4 + Fold Change Top3 | 0.95 | 57.41 | -14.32 |
| Model 11 + Creatinine |  | 0.96 | 57.74 | -14.86 |

### Refining predictors of DKD progression by patient characteristics matching

One caveat of utilizing all 75 patients for multivariate model development is that serum creatinine concentrations already differ significantly between the DKD progressors and non-progressors at the time of enrollment. Thus, the identified biomarkers may be consequences of discrepancies in serum creatinine content rather than predictors independent of baseline serum creatinine [39]. To identify variables independent of serum creatinine concentration, patients with high and low baseline serum creatinine concentrations were excluded. This enabled DKD progressors and non-progressors exhibit approximately equivalent baseline serum creatinine. Multivariate analysis was then performed. Combining the results of univariate and multivariate analyses, with or without the inclusion of baseline serum creatinine as a variate, 10 models were evaluated (**TABLE 4**). The AIC value was then used to identify the optimal model (model 7, AUC Top4 + Fold Change Top2) as the best one, with an AIC value of 49.50 and an AUC value of 0.91. Notably, we found 2 models that achieved AUC values of about 0.90 (**FIGURE 10, A-C**). Consistent with studies using all 75 samples, inclusion of baseline serum creatinine as a variate did not improve these models (**FIGURE 10, D-F**).

**TABLE 4.** Multivariate model selection after patient serum creatinine concentration matching. After removing samples with high and low baseline serum creatinine values, starting serum creatinine concentration was similar in P and NP groups. Performance of all multivariate models was assessed, along with top models (AUC > 0.90) with addition of serum creatinine as a variate. Model performance was compared based on AUC, AIC and intercept values.

| Group | Combination Name | AUC | AIC | Intercept |
| --- | --- | --- | --- | --- |
| Model 1 | Random Forest Top10 | 0.90 | 62.62 | -0.82 |
| Model 1 + Creatinine |  | 0.90 | 64.56 | -0.25 |
| Model 2 | LASSO Top3 | 0.65 | 77.08 | 2.08 |
| Model 3 | Random Forest Top3 | 0.80 | 64.03 | 12.45 |
| Model 4 | Random Forest Top3 + LASSO Top3 | 0.83 | 64.85 | 21.13 |
| Model 5 | Random Forest Top2 + LASSO Top2 | 0.81 | 63.82 | 11.76 |
| Model 6 | Random Forest Top1 + LASSO Top1 | 0.70 | 71.76 | 6.42 |
| Model 7 | AUC Top4 + Fold Change Top2 | 0.91 | 49.50 | -21.44 |
| Model 7 + Creatinine |  | 0.91 | 51.07 | -25.77 |
| Model 8 | AUC Top2 + Fold Change Top2 | 0.87 | 52.54 | 2.77 |

**FIGURE 10.**
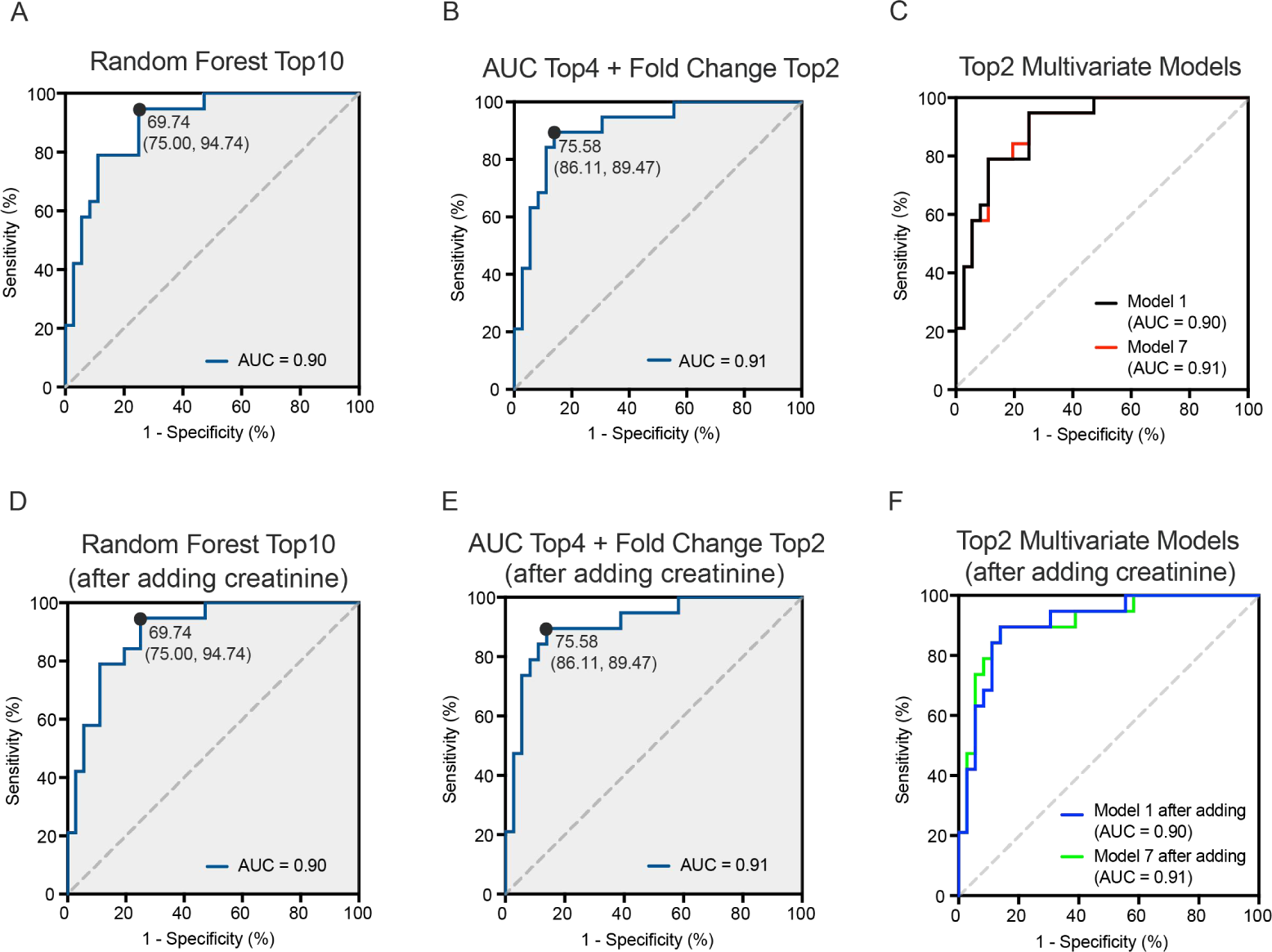
Assessment of top multivariate models after matching baseline serum creatinine. (A-B) ROC curves of top 2 multivariate models. AUC value, Youden index (sensitivity, specificity) are indicated in each panel. (C) ROC curves of top multivariate models. (D-E) ROC curves of top 2 models with addition of baseline serum creatinine as a variate. AUC value, Youden index (sensitivity, specificity) are indicated in each panel. (F) ROC curves of top 2 multivariate models after the addition of baseline serum creatinine as a variable.

### Properties of lead predictors

Several candidate proteins were identified by multiple feature selection methods and as variates contributing to the discriminative power of leading multivariate models. Our analysis showed that prostaglandin-H2 D-isomerase (PTGDS, P41222) is a leading candidate differentiating DKD progressors and non-progressors. This enzyme facilitates in isomerization of prostaglandin H2 to prostaglandin D2 [40]. PTGDS is has been shown to be highly expressed in tumor endothelial cells, and modulates the biosynthesis of PGD2, which restrains aberrant proliferation and migration of tumor endothelial cells and ultimately attenuating tumor angiogenesis [41]. Another designation for Prostaglandin-H2 D-isomerase is β-trace protein (BTP), and as the name implies, it can be filtered through glomerulus and primarily reabsorbed in renal tubules. Its concentration exhibits an inverse correlation with the glomerular filtration rate (GFR). Elevated BTP expression might also display a positive association with renal tubular injury and serve as a biomarker for kidney function and cardiovascular disease [42].

Von Willebrand factor (VWF, P04275) is a large, multi-domain glycoprotein primarily synthesized by endothelial cells and megakaryocytes and subsequently released into the plasma [43]. VWF plays a pivotal role in primary hemostasis and platelet aggregation [44]. Research has found that patients with end-stage renal disease exhibit elevated plasma VWF levels, and the functional activity of VWF is reduced. These changes lead to an increased risk of mortality in patients [45].

Band 3 anion transport protein (B3AT, P02730) is a transport protein located on the cell membrane that mediates electroneutral anion exchange. It mediates 1:1 exchange of chloride and bicarbonate exchange, in red blood cell membrane and kidney epithelial cells [46–48]. B3AT protein is coded by the *SLC4A1*gene. Patients with *SLC4A1*gene mutations have been identified, and these patients exhibit failure of the α-intercalated cells in the distal renal tubular cortical collecting duct, resulting in reduced urine acidification, variable hyperchloremic hypokalemic metabolic acidosis, nephrocalcinosis, and kidney stones, causing renal tubular acidosis [49–51].

Beta-2-microglobulin (B2MG, P61769) is a low molecular weight protein. B2MG can serve as an early predictive biomarker for acute kidney injury in neonates with perinatal asphyxia [52]. It can also be combined with BTP to form a new GFR estimating equation that outperforms creatinine-based equations, especially in populations with impaired kidney function and the elderlies [53].

## DISCCUSSION

In this prospective study, we have developed a Proteonano™ platform for non-targeted deep proteomics to identify serum protein biomarkers that differentiate DKD patients that progressed in up to 5 years of follow up. A total of 75 patients of East Asian descent were serially enrolled and serum samples were prepared at the time of recruitment. Using these samples, we have identified a set of proteins that are differentially expressed in DKD patients that progressed compared to those who did not progress, and established combination of serum protein markers for sensitive and specific identification of DKD patients with high likelihood of progression.

We have developed a nanobinder-based proteomic technology that has shown the ability to consistently detect more than 1,000 plasma proteins by label free quantification. In this cohort, we detected a total of 1,393 proteins, and 469 proteins were commonly detected in all samples. Using relative quantification, 38 significantly different proteins were identified, and about 20 of these serum proteins have AUC number similar to that of serum creatinine concentration. Combinatory analysis of detected proteins allowed us to identify algorithms that have AUC values higher than 0.95. Noticeably, adding creatinine to these models had little impact on AUC values, indicating these serum protein biomarkers can efficiently distinct DKD progressors and non-progressors. Furthermore, when serum creatinine is included in random forest and LASSO models, the resultant models have lower AUC values, indicating serum creatinine is likely an inferior predictor than combination of serum protein markers described here.

The patients enrolled have significantly different serum creatinine levels, which alone could drive significantly different serum protein compositions. Thus, we removed patients with high and low baseline serum creatinine levels. Although this reduced the number of patients included in analysis from 75 to 55 decreased statistical power, it allowed baseline serum creatinine between progressors and non-progressors to be matched. While only a small number of proteins remained to have borderline protein abundance difference, combining random forest, LASSO, protein abundance difference and univariate ROC analyses was still able to identify multivariate models that have AUC numbers higher than 0.90. Furthermore, including creatinine in analysis did not improve these models. This further suggests that proteins identified in this deep proteomic analysis was able to identify promising potential prognostic biomarkers for DKD progression.

Taking together, these results demonstrate that our Proteonano™ Platform for untargeted deep proteomics allowed us to develop excellent prognostic models that can distinguish DKD progressors and non-progressors with high sensitivity and specificity.

Despite these advances, our current study has its own limitations and many following works to do before it can be translated into a multi-protein test kit used in clinical. First, the clinical samples used in this study are stored in −80 °C for at least 5 years. Fresher plasma samples with proper quality control methods would be desirable for further study. Second, only a relatively small number of patients was used in this study, and including more patients in future studies may allow us to identify additional multivariate models with better sensitivity and specificity. Second, the models generated in this discovery cohort need to be validated. This may involve using absolute quantification methods with more samples enrolled for further analyses. Such methods include targeted proteomics, and antibody-based protein quantification. The most discriminative model created in our analysis was top 10 markers identified by random forest analysis, with an AUC of 0.97. This combination is unlikely to be adopted for clinical use, due to the large number of proteins that need to be tested. Our exploration of other models with reduced number of proteins allowed us constructing several models with only slightly reduced sensitivity and specificity, and these models can be further optimized.

Despite these caveats, our prognostic models incorporated a number of proteins that have not been implicated previously in DKD. Among the proteins identified, prostaglandin D2 synthase (PTGDS, encoded by *PTGDS*gene), Von Willebrand factor (VWF, encoded by *VWF*gene), and band 3 anion transport protein (B3AT, encoded by *SLC4A1*gene) are three of the most often presented proteins in these models. PTGDS is an enzyme that converts PGH2 to PGD2, and PGD2 regulates smooth muscle cell contraction and relaxation, and is an important inhibitor for platelet aggregation, which may regulate vascular function during endothelial damage that occurs in DKD [54]. Although PTGDS itself has not been implicated in loss of kidney function, it has an important paralog, lipocalin 2 (LCN2) that is being considered as a biomarker for an early loss of kidney function [55–57]. VWF is a well-recognized factor for hemostasis, and is considered a marker for endothelial cells [58, 59]. It regulates many endothelial related processes, including platelet aggregation at sites of endothelial damage [59]. Indeed, high plasma VWF has previously been implicated with decreased kidney function and development of microalbuminuria in both diabetic and non-diabetic patients [58–60]. B3AT is an anion exchanger that regulates HCO3^−^ and Cl^−^ transport [61, 62]. Besides its expression in erythrocytes, it is also expressed in distal renal tubules [62]. Patients with mutated the B3AT coding gene *SLC4A1*have been identified, and some of these patients have defective kidney acid secretion, leading to distal renal tubular acidosis [61–63]. Thus, proteins identified in this study that can be used as biomarkers for DKD may also participate in DKD pathogenesis and progression. Further studies are needed to establish mechanistic links between these proteins and DKD, which may also lead to discovery of novel targets for DKD treatment.

In addition to identifying novel biomarkers for DKD progression, our study further validated Proteonano™ Platform as a great sample preprocessing method to increase the depth of proteomics-based biomarker discovery. Further optimization of nanobinders for protein enrichment and utilization of an expanded panel of nanobinders that can profile distinctive groups of proteins may have broad implications in promoting biomarker discovery.

In conclusion, our Proteonano™ Platform shows its excellent capacity in deep proteomics study, with a pg/mL sensitivity and 1000-plex, which enables us to identify potentially new biomarkers for DKD progression and to construct prognostic models to identify patients at risk for DKD progression with high accuracy and specificity. Further studies leveraging these findings will advance our understanding of molecular and cellular mechanisms for DKD initiation and progression, and will allow us developing diagnostic tools that identify patients that are likely to have DKD progression. These will eventually lead to significantly improved care for DKD patients.

## METHODS

### Patient recruitment and follow up

75 patients with primary clinical diagnosis of diabetic kidney disease, but not other kidney disease and hypertension were serially enrolled from Beijing Hospital nephrology clinic between 2012 and 2019 with ethical committee approval. Patients were followed up to five years. Clinical parameters of these patients were collected from medical records, and stored in deidentified manner for data analysis.

### Blood collection and serum sample preparation

Blood samples were collected at Beijing Hospital Nephrology Clinic by phlebotomists. Whole blood was incubated in serum tubes and allowed to clot at ambient temperature. Samples were centrifuged 30 min later at 1500 *g* for 15 min. Supernatant was transferred to microfuge tubes, aliquoted, and stored in −80 °C until analysis.

At the time of enrollment and during follow up, clinical test samples were also collected. Serum creatinine was determined at Department of Laboratory Medicine, Beijing Hospital as part of regular clinical care.

### Proteonano based serum sample processing for proteomics

Semi-automatic sample preparation was performed in discrete batches utilizing an OT-2 automated workstation (Opentrons, New York, NY, USA), equipped with both a magnetic module and a heater shaker module (Opentrons, New York, NY, USA). Each batch was composed of 16 samples, including 15 serum samples and a human plasma standard sample for quality control. 20 μL of serum specimens were diluted to a final volume of 100 μL by 1x PBS and were subsequently combined with nanobinders from the Proteonano™ proteomic assay kit (Nanomics Biotechnology Co., Ltd., Hangzhou, Zhejiang, China) in a 1:1 volumetric ratio. The mixture was incubated at 25 °C and agitated at a rate of 1700 rpm for a duration of 60 min. After magnetic immobilization, nanobinders were washed thrice with 1 x PBS. Proteins captured on the nanobinders were reduced by 20 mM DTT at 37 °C for 60 min. Next, alkylation was performed using 50 mM IAA at room temperature in complete darkness for 30 min. Subsequent to these steps, trypsin digestion (V5111, Promega Corporation, Madison, WI, USA) was carried out at 37 °C for a duration of 16 hours with shaking at 1000 rpm. Post-digestion, peptides were purified using a C18 desalting column (87784, Thermo Fisher Scientific, Waltham, MA, USA) and lyophilized with a LyoQuest freeze dryer (LyoQuest, Telstar, Terrassa, Spain). Lyophilized peptides were then reconstituted in 0.1 % formic acid prior to mass spectrometry. Peptide concentrations were measured with a Nano300 microvolume spectrophotometer (Aosheng Instruments, China), and 500 *ng*of peptide from each sample was subjected to LC-MS/MS.

### Label free LC-MS analysis

Peptides were separated by an Easy-nLC1200 reverse-phase HPLC system (Thermo Fisher, Waltham, MA) using a precolumn (made in house, 0.075 mm × 2 cm, 1.9 μm, C18) and a self-packed analytical column (0.075 mm × 20 cm, 1.9 μm, C18) over a 48 min gradient before nano-electrospray on Orbitrap Exploris 480 mass spectrometer equipped with FAIMS (Thermo Fisher, Waltham, MA). Solvent A was 0.1 % formic acid and solvent B was 80 % ACN/0.1 % formic acid. Gradient conditions were 3–7 % B (0–1 min), 7–30 % B (1–36 min), 30–95 % B (36–38 min), 95 % B (38–48 min). Mass spectrometer was operated in data-independent mode (DIA). The spray voltage was set to 2.4 kV, RF Lens level at 40 %, and heated capillary temperature at 320 °C. For DIA experiments, full MS resolutions were set to 60,000 at m/z 200 and full MS automatic gain control (AGC) target was 100 % with an injection time (IT) of 50 ms. Mass range was set to m/z 350–1200. The AGC target value for fragment spectra was set at 1,000 %. Resolution was set to 30,000 and IT to 54 ms. Normalized collision energy was set at 30 %. Default settings were used for FAIMS with voltages applied as −45 V, except gas flow, which was applied with 3.5 L/min.

### Proteomics data processing

Raw mass spectral data files (.raw) were converted to .mzML by MSConvert GUI (Vendoer, Version 3.0) [64]. All .mzML files were searched using DIA-NN software (version 1.8.1) [64] in library free mode to generate a spectral library. This library was searched against a protein library containing 20,422 Entries (UniProt *Homo sapiens* reviewed proteome dataset, UP000005640). DIA-NN search parameters were: 10 ppm mass tolerance for mass accuracy, 1 missed cleavages of trypsin, carbamidomethylation of cysteine as fixed modification, and methionine oxidation as the only variable modification. The rest of the parameters were set to default. The FDR cutoffs at both precursor and protein level were set to 0.01.

Data generated by DIA-NN software were imported into Perseus (version 2.0.10.0) [65] for processing, following a specific workflow as outlined here. Intensity values were log_2_ transformed, followed by data standardization with the median of z-scores. Data were then categorized into non-progressors (NP) and progressors (P) and a filtering criterion was applied at least one group to contain 70 % of the data. Remaining missing values were imputed by drawing random samples from a normal distribution with downshifted mean by 1.8 SD and scaled SD (0.3) relative to that of abundance distribution of all proteins in one sample. Finally, two-sample test with a significance level (p-value) of 0.05 and s0 = 0 was employed to compare P and NP groups, resulting in the identification of 38 significantly differentially expressed proteins.

### Bioinformatic analyses

Data generated by the DIA-NN software was first analyzed from an overall perspective, utilizing the ggplot2 in R to visualize the total protein quantities in each group as boxplots and violin plots [66]. Following this, Venn diagrams were created by using TBtools [67]. GO biological process and KEGG pathway enrichment analysis was performed using the R clusterProfiler software package(Bioconductor - clusterProfiler) [68]. The critical value of FDR (adj. p) < 0.05 was considered statistically significant. Functional associations between proteins were predicted by using the STRING version 12.0 online tool with medium-confidence association score of ≥ 0.4 [69].

### Biostatistics analyses

To determine the predictive power of 38 candidate proteins for DKD progression, univariate analysis was performed. Box plots and receiver operating characteristic (ROC) curves were graphed using Prism 9.0 (GraphPad, Boston, MA). Odds ratio was determined by using ggplot2 in R (version 4.0.3, https://www.r-project.org/) [64]. For correlation analysis, Pearson correlation heatmaps were generated using Prism 9.0. Differential fold change (FC) analysis was carried out in MetaboAnalyst 5.0 with a threshold of 2.0 (|log_2_FC| > 0.5) [70]. Feature selection was performed using random forest (randomForest in R, through MetaboAnalyst), and the least absolute shrinkage and selection operator (LASSO, glmnet package in R) methods [70, 71]. Multivariate analysis was performed using MASS in R [72]. Variables were first processed by glm (MASS package in R), followed by Akaike information criterion (AIC) determination using stepAIC (MASS package in R) with forward selection and Wald test. Multivariate ROC analysis was subsequently performed using roc (glmtoolbox package in R) to assess the predictive performance of the logistic regression model selected by stepAIC [73].

## ACKNOWLEDGEMENTS

This work was supported by grants from the Beijing Hospital (IIT-2021-09-03, 121-2016008).

## DISCLOSURE

X.C.G., X.H.O., J.K.F., Z.H.D and H.W. are employees of Nanomics Biotechnology Co., Ltd. Other authors declare no competing interests.

## AUTHOR CONTRIBUTIONS

B.Z., H.W. and Y.H.M. conceived and designed the study. B.Z., X.H.O., J.K.F and Z.H.D. performed experiments. B.Z., X.C.G, J.K.F and Y.H.M. analyzed and interpreted the data. X.C.G, J.K.F, X.H.O, and H.W. drafted the manuscript. All authors have reviewed and approved the final manuscript.

## REFERENCES

1. Chen, L., D.J. Magliano, and P.Z. Zimmet, The worldwide epidemiology of type 2 diabetes mellitus—present and future perspectives. Nature reviews endocrinology, 2012. 8(4): p. 228–236.

2. Thomas, M.C., et al., Diabetic kidney disease. Nature reviews Disease primers, 2015. 1(1): p. 1–20.

3. Reidy, K., et al., Molecular mechanisms of diabetic kidney disease. The Journal of clinical investigation, 2014. 124(6): p. 2333–2340.

4. Barutta, F., et al., Novel biomarkers of diabetic kidney disease: current status and potential clinical application. Acta Diabetologica, 2021. 58: p. 819–830.

5. White, C.A., et al., Estimating glomerular filtration rate in kidney transplantation: is the new chronic kidney disease epidemiology collaboration equation any better? Clinical chemistry, 2010. 56(3): p. 474–477.

6. Rico-Fontalvo, J., et al., Novel Biomarkers of Diabetic Kidney Disease. Biomolecules, 2023. 13(4): p. 633.

7. Colhoun, H.M. and M.L. Marcovecchio, Biomarkers of diabetic kidney disease. Diabetologia, 2018. 61(5): p. 996–1011.

8. Jung, C.-Y. and T.-H. Yoo, Pathophysiologic mechanisms and potential biomarkers in diabetic kidney disease. Diabetes & Metabolism Journal, 2022. 46(2): p. 181–197.

9. Good, D.M., et al., Naturally occurring human urinary peptides for use in diagnosis of chronic kidney disease. Molecular & cellular proteomics, 2010. 9(11): p. 2424–2437.

10. Rodríguez-Ortiz, M.E., et al., Novel urinary biomarkers for improved prediction of progressive eGFR loss in early chronic kidney disease stages and in high risk individuals without chronic kidney disease. Scientific reports, 2018. 8(1): p. 15940.

11. Lindhardt, M., et al., Urinary proteomics predict onset of microalbuminuria in normoalbuminuric type 2 diabetic patients, a sub-study of the DIRECT-Protect 2 study. Nephrology Dialysis Transplantation, 2017. 32(11): p. 1866–1873.

12. Gold, L., et al., Aptamer-based multiplexed proteomic technology for biomarker discovery. Nature Precedings, 2010: p. 1–1.

13. Nishi, H., Aptamer-Based Proteomic Platform for Human Immune-Mediated Kidney Diseases. 2022, Elsevier. p. 1450–1452.

14. Wu, P.-H., et al., Novel biomarkers detected by proteomics predict death and cardiovascular events in hemodialysis patients. Biomedicines, 2022. 10(4): p. 740.

15. Petrackova, A., et al., Serum protein pattern associated with organ damage and lupus nephritis in systemic lupus erythematosus revealed by PEA immunoassay. Clinical proteomics, 2017. 14(1): p. 1–15.

16. Zubiri, I., et al., Diabetic nephropathy induces changes in the proteome of human urinary exosomes as revealed by label-free comparative analysis. Journal of proteomics, 2014. 96: p. 92–102.

17. Fu, J., et al., Advances in current diabetes proteomics: from the perspectives of label-free quantification and biomarker selection. Current Drug Targets, 2020. 21(1): p. 34–54.

18. Poleti, M.D., et al., Longissimus dorsi muscle label-free quantitative proteomic reveals biological mechanisms associated with intramuscular fat deposition. Journal of proteomics, 2018. 179: p. 30–41.

19. Distler, U., et al., Drift time-specific collision energies enable deep-coverage data-independent acquisition proteomics. Nature methods, 2014. 11(2): p. 167–170.

20. Zhao, L., et al., Comparative proteomics reveals genetic mechanisms of body weight in Hu sheep and Dorper sheep. Journal of Proteomics, 2022. 267: p. 104699.

21. Schwenk, J.M., et al., The human plasma proteome draft of 2017: building on the human plasma PeptideAtlas from mass spectrometry and complementary assays. Journal of proteome research, 2017. 16(12): p. 4299–4310.

22. Anderson, N.L. and N.G. Anderson, The human plasma proteome: history, character, and diagnostic prospects. Molecular & cellular proteomics, 2002. 1(11): p. 845–867.

23. Looker, H.C., et al., Protein biomarkers for the prediction of cardiovascular disease in type 2 diabetes. Diabetologia, 2015. 58: p. 1363–1371.

24. Liu, D., et al., Identification of ferroptosis-related genes and pathways in diabetic kidney disease using bioinformatics analysis. Scientific Reports, 2022. 12(1): p. 22613.

25. Bonnans, C., J. Chou, and Z. Werb, Remodelling the extracellular matrix in development and disease. Nature reviews Molecular cell biology, 2014. 15(12): p. 786–801.

26. Ruggeri, Z.M., Platelets in atherothrombosis. Nature medicine, 2002. 8(11): p. 1227–1234.

27. Sira, J. and L. Eyre, Physiology of haemostasis. Anaesthesia & Intensive Care Medicine, 2016. 17(2): p. 79–82.

28. Heijnen, H. and P. Van Der Sluijs, Platelet secretory behaviour: as diverse as the granules… or not? Journal of thrombosis and haemostasis, 2015. 13(12): p. 2141–2151.

29. Rabouille, C. and G. Haase, Golgi pathology in neurodegenerative diseases. 2016, Frontiers Media SA. p. 489.

30. Shavit-Stein, E., et al., The role of thrombin in the pathogenesis of diabetic neuropathy. PLoS One, 2019. 14(7): p. e0219453.

31. Szklarczyk, D., et al., The STRING database in 2017: quality-controlled protein–protein association networks, made broadly accessible. Nucleic acids research, 2016: p. gkw937.

32. Widner-Andrä, R.A., Assignment of functional impact on genetic data in two mouse models of affective disorders. 2012, lmu.

33. Shao, B., et al., A cluster of proteins implicated in kidney disease is increased in high-density lipoprotein isolated from hemodialysis subjects. Journal of proteome research, 2015. 14(7): p. 2792–2806.

34. Klingenstein, A., et al., Receiver operating characteristic analysis: calculation for the marker ‘melanoma inhibitory activity’in metastatic uveal melanoma patients. Melanoma Research, 2011. 21(4): p. 352–356.

35. Kukreja, S.L., J. Löfberg, and M.J. Brenner, A least absolute shrinkage and selection operator (LASSO) for nonlinear system identification. IFAC proceedings volumes, 2006. 39(1): p. 814–819.

36. Kursa, M.B. and W.R. Rudnicki, Feature selection with the Boruta package. Journal of statistical software, 2010. 36: p. 1–13.

37. Burnham, K.P. and D.R. Anderson, Multimodel inference: understanding AIC and BIC in model selection. Sociological methods & research, 2004. 33(2): p. 261–304.

38. Doshi, S.M. and A.N. Friedman, Diagnosis and management of type 2 diabetic kidney disease. Clinical journal of the American Society of Nephrology: CJASN, 2017. 12(8): p. 1366.

39. Jylhä, M., S. Volpato, and J.M. Guralnik, Self-rated health showed a graded association with frequently used biomarkers in a large population sample. Journal of clinical epidemiology, 2006. 59(5): p. 465–471.

40. Urade, Y. and O. Hayaishi, Prostaglandin D synthase: structure and function. 2000.

41. Omori, K., et al., Lipocalin-type prostaglandin D synthase-derived PGD2 attenuates malignant properties of tumor endothelial cells. The Journal of pathology, 2018. 244(1): p. 84–96.

42. White, C.A., S. Ghazan-Shahi, and M.A. Adams, β-Trace protein: a marker of GFR and other biological pathways. American Journal of Kidney Diseases, 2015. 65(1): p. 131–146.

43. Wagner, D.D., Cell biology of von Willebrand factor. Annual review of cell biology, 1990. 6(1): p. 217–242.

44. Ruggeri, Z., Von Willebrand factor, platelets and endothelial cell interactions. Journal of thrombosis and haemostasis, 2003. 1(7): p. 1335–1342.

45. Holden, R.M., et al., Quantitative and qualitative changes of von Willebrand factor and their impact on mortality in patients with end-stage kidney disease. Blood Coagulation & Fibrinolysis, 2013. 24(7): p. 719–726.

46. Rungroj, N., et al., A novel missense mutation in AE1 causing autosomal dominant distal renal tubular acidosis retains normal transport function but is mistargeted in polarized epithelial cells. Journal of Biological Chemistry, 2004. 279(14): p. 13833–13838.

47. Bruce, L.J., et al., Monovalent cation leaks in human red cells caused by single amino-acid substitutions in the transport domain of the band 3 chloride-bicarbonate exchanger, AE1. Nature genetics, 2005. 37(11): p. 1258–1263.

48. Shnitsar, V., et al., A substrate access tunnel in the cytosolic domain is not an essential feature of the solute carrier 4 (SLC4) family of bicarbonate transporters. Journal of Biological Chemistry, 2013. 288(47): p. 33848–33860.

49. Bruce, L.J., et al., Band 3 mutations, renal tubular acidosis and South-East Asian ovalocytosis in Malaysia and Papua New Guinea: loss of up to 95% band 3 transport in red cells. Biochemical journal, 2000. 350(1): p. 41–51.

50. Sritippayawan, S., et al., Novel compound heterozygous SLC4A1 mutations in Thai patients with autosomal recessive distal renal tubular acidosis. American journal of kidney diseases, 2004. 44(1): p. 64–70.

51. Yenchitsomanus, P.-t., et al., Anion exchanger 1 mutations associated with distal renal tubular acidosis in the Thai population. Journal of human genetics, 2003. 48(9): p. 451–456.

52. Abdullah, et al., Urinary beta-2 microglobulin as an early predictive biomarker of acute kidney injury in neonates with perinatal asphyxia. European Journal of Pediatrics, 2022. 181(1): p. 281–286.

53. Inker, L.A., et al., GFR estimation using β-trace protein and β2-microglobulin in CKD. American journal of kidney diseases, 2016. 67(1): p. 40–48.

54. Takahashi, N., et al., Deletion of Alox15 improves kidney dysfunction and inhibits fibrosis by increased PGD 2 in the kidney. Clinical and experimental nephrology, 2021. 25: p. 445–455.

55. Perco, P., et al., Protein biomarkers associated with acute renal failure and chronic kidney disease. European journal of clinical investigation, 2006. 36(11): p. 753–763.

56. Viau, A., et al., Lipocalin 2 is essential for chronic kidney disease progression in mice and humans. The Journal of clinical investigation, 2010. 120(11): p. 4065–4076.

57. Nickolas, T.L., et al., NGAL (Lcn2) monomer is associated with tubulointerstitial damage in chronic kidney disease. Kidney international, 2012. 82(6): p. 718–722.

58. van der Vorm, L.N., et al., Circulating active von Willebrand factor levels are increased in chronic kidney disease and end-stage renal disease. Clinical kidney journal, 2020. 13(1): p. 72–74.

59. Shen, L., et al., Von Willebrand factor, ADAMTS13 activity, TNF-α and their relationships in patients with chronic kidney disease. Experimental and therapeutic medicine, 2012. 3(3): p. 530–534.

60. Hirano, T., et al., Vascular endothelial markers, von Willebrand factor and thrombomodulin index, are specifically elevated in type 2 diabetic patients with nephropathy: comparison of primary renal disease. Clinica chimica acta, 2000. 299(1-2): p. 65–75.

61. Shayakul, C., et al., Characterization of a highly polymorphic marker adjacent to the SLC4A1 gene and of kidney immunostaining in a family with distal renal tubular acidosis. Nephrology Dialysis Transplantation, 2004. 19(2): p. 371–379.

62. Vichot, A.A., et al., Loss of kAE1 expression in collecting ducts of end-stage kidneys from a family with SLC4A1 G609R-associated distal renal tubular acidosis. Clinical Kidney Journal, 2017. 10(1): p. 135–140.

63. Mohebbi, N. and C.A. Wagner, Pathophysiology, diagnosis and treatment of inherited distal renal tubular acidosis. Journal of nephrology, 2018. 31(4): p. 511–522.

64. Schratz, P. and M.P. Schratz, Package ‘oddsratio’. 2020.

65. Tyanova, S. and J. Cox, Perseus: a bioinformatics platform for integrative analysis of proteomics data in cancer research. Cancer systems biology: Methods and protocols, 2018: p. 133–148.

66. Gómez-Rubio, V., ggplot2-elegant graphics for data analysis. Journal of Statistical Software, 2017. 77: p. 1–3.

67. Chen, C., et al., TBtools: an integrative toolkit developed for interactive analyses of big biological data. Molecular plant, 2020. 13(8): p. 1194–1202.

68. Yu, G., et al., clusterProfiler: an R package for comparing biological themes among gene clusters. Omics: a journal of integrative biology, 2012. 16(5): p. 284–287.

69. Szklarczyk, D., et al., The STRING database in 2023: protein–protein association networks and functional enrichment analyses for any sequenced genome of interest. Nucleic acids research, 2023. 51(D1): p. D638–D646.

70. Pang, Z., et al., MetaboAnalyst 5.0: narrowing the gap between raw spectra and functional insights. Nucleic acids research, 2021. 49(W1): p. W388–W396.

71. Fonti, V. and E. Belitser, Feature selection using lasso. VU Amsterdam research paper in business analytics, 2017. 30: p. 1–25.

72. Zhang, Z., Variable selection with stepwise and best subset approaches. Annals of translational medicine, 2016. 4(7).

73. Vanegas, L., L. Rondón, and G. Paula, glmtoolbox: Set of tools to data analysis using generalized linear models. Preprint at, 2022.

